# Topological analysis of a bacterial DedA protein associated with alkaline tolerance and antimicrobial resistance

**DOI:** 10.1101/2021.10.18.464800

**Authors:** Hollie L Scarsbrook, Roman Urban, Bree R. Streather, Alexandra Moores, Christopher Mulligan

## Abstract

Maintaining membrane integrity is of paramount importance to the survival of bacteria as the membrane is the site of multiple crucial cellular processes including energy generation, nutrient uptake, and antimicrobial efflux. The DedA family of integral membrane proteins are widespread in bacteria and are associated with maintaining the integrity of the membrane. In addition, DedA proteins have been linked to resistance to multiple classes of antimicrobials in various microorganisms. Therefore, the DedA family are attractive targets for the development of new antibiotics. Despite DedA family members playing a key physiological role in many bacteria, their structure, function and physiological role remain unclear. To help illuminate the structure of the bacterial DedA proteins, we have performed substituted cysteine accessibility method (SCAM) analysis on the most comprehensively characterized bacterial DedA protein, YqjA from *Escherichia coli*. By probing the accessibility of 15 cysteine residues across the length of YqjA using thiol reactive reagents, we have mapped the topology of the protein. Using these data, we have experimentally validated a structural model of YqjA generated using evolutionary co-variance, which consists of an α-helical bundle with two re-entrant hairpin loops reminiscent of several secondary active transporters. In addition, our cysteine accessibility data suggests that YqjA forms an oligomer wherein the protomers are arranged in a parallel fashion. This experimentally verified model of YqjA lays the foundation for future work in understanding the function and mechanism of this interesting and important family.

## Introduction

The bacterial cell envelope is involved in many crucial processes from interaction with the environment, signaling and nutrient uptake to energy production. As such, maintaining the integrity of the membrane is of paramount importance to the survival of the bacterium during the various rigours of its existence. Therefore, targeting proteins involved in the maintenance of membrane integrity has great potential for the production or enhancement of antimicrobials. One such protein family is the DedA superfamily of integral membrane proteins that are found widespread in bacteria, including many human pathogens^1,2^. Members of the DedA superfamily are enigmatic; their function and physiological role remain unclear. However, the effects of disrupting DedA function are substantial. The *E. coli* genome encodes 8 DedA genes and they are collectively essential^3^; in addition, the single DedA gene encoded by the *Borrelia burgdorferi* genome is also essential^4^, indicating a crucial physiological role for this protein family. Deletion of the genes encoding two DedA family members in *E. coli*, YqjA and YghB, results in a pleiotropic phenotype including sensitivity to elevated temperatures and pH^5–8^, cell division defects caused by an inability to secrete periplasmic amidases^9^; and sensitivity to multiple antimicrobial agents^10^. This sensitivity to antimicrobial compounds can be mitigated by reducing the pH of the growth medium thus increasing the proton motive force, increasing the extracellular Na^+^ concentration, or by overexpression of *mdfA*, which encodes a proton-coupled multidrug efflux transporter, which also moonlights as a monovalent cation/H^+^ exchanger^10^. These findings, and the observation that two conserved, membrane embedded acidic residues are essential for function^10^, strongly suggests that members of the DedA superfamily have transport activity likely involving proton flux. Beyond *E. coli*, members of the DedA superfamily are required for colistin resistance in *Klebsiella pneumoniae* and *Burkholderia thailandensis*^11–13^, and are involved in resistance to cationic antimicrobial peptides (CAMP) in *Salmonella enterica* and *Neisseria meningitidis*^14–15^, and the macrocyclic alkaloid, halicyclamine A in *Mycobacterium bovis*^16^, further highlighting the broad potential impact of understanding DedA structure and function for the treatment of drug-resistant infections.

Understanding the function and mechanism of the DedA family has been hampered by a lack of experimentally derived structural information. Based on hydropathy profile alignments it has been proposed that the DedA family share a similar fold to the LeuT family of transporters^17^. LeuT transporters consist of an inverted structural repeat consisting of 5 transmembrane helices that evolved via gene duplication and inversion of a 5 TM progenitor^18^. Due to the similarity of their hydropathy profiles which reports on general structural features, the DedA superfamily was proposed to be the 5 TM LeuT progenitor, suggesting that DedA proteins form dual topology oligomers in the membrane^17^. More recently, evolutionary co-variance analysis has been used to generate a 3D model for members of the DedA superfamily, suggesting the presence of 2 re-entrant hairpin loops^2,19^. While this topology has been partially experimentally verified for a human DedA superfamily member^2^, there has been no experimental validation of the structural arrangement of any bacterial DedA protein, which are distantly related to the human homologues and may have diverged in function^1^.

Here, we have sought to gain a better understanding of the structure of bacterial DedA proteins by performing substituted cysteine accessibility method (SCAM) on the most comprehensively characterized DedA protein, YqjA, from *E. coli*. Our data reveal that the accessibility of several residues does not match what is expected based on topology model prediction software that is traditionally used to investigate membrane protein topology. However, our SCAM data do support a structural model generated using evolutionary co-variance analysis, which, in support of previous studies^2,19^, predicts a compact structure for YqjA comprised of 2 re-entrant hairpins. Furthermore, our data strongly indicate that YqjA forms an oligomer in which the subunits are arranged in a strictly parallel fashion.

## Methods

### Generation of the E. coli BW25113ΔΔ strain

*Escherichia coli* strains used during this study are derivatives of Keio Collection BW25113 K-12^20^. The BW25113ΔyqjAΔyghB (BW25113ΔΔ) double mutant was created by first eliminating the Kanamycin cassette from JW3066-1 (BW25113ΔyqjA) using site-specific FLP recombination via pCP20^21^. Subsequently, the BW25113ΔyghB mutation was transferred from JW2976-2 strain through P1 transduction carried out using P1_vir_ bacteriophage^22^. Both mutations were and verified by PCR/sequencing.

### Site directed mutagenesis

All mutagenesis was performed using the QuikChange II Site-directed mutagenesis kit (Agilent). The accuracy of all constructs was checked by sequencing.

### Phenotypic rescue assays

To perform the temperature sensitivity rescue assay, BW25113wt or BW25113ΔΔ cells were freshly transformed with arabinose-inducible pBAD-based plasmid encoding a variant of *yqjA* or *gltph* (a non-DedA membrane protein to be used to control for the effects overexpression of membrane proteins has on bacterial growth). The transformed strains were grown overnight in LB supplemented with 100 μg/ml ampicillin, harvested, normalized to an OD_600_ of 1 using, then serially diluted. 5 μl of each dilution was spotted onto LB agar supplemented with 100 μg/ml ampicillin and 0.001% (w/v) L-arabinose. Once dry, the plates were incubated at a permissive temperature of 30°C or non-permissive temperature of 44°C for ~16 h.

### Substituted cysteine accessibility method (SCAM) assay

To perform SCAM, *E. coli* TOP10 were freshly transformed with pBAD plasmid encoding a variant of *yqjA* upstream of a histidine tag. A single colony of transformed cells was grown overnight at 37°C in LB supplemented with 100 μg/ml ampicillin. Following overnight incubation, the cultures were diluted to an OD_600_ of 0.2 using fresh LB supplemented with ampicillin and grown at 30°C for 1.5 h. Protein expression was then induced by addition of 0.1% (w/v) L-arabinose and the cells were grown for 1 h. Cells were harvested, resuspended in PBS to an OD between 0.6-0.8 then divided into 4 equal volume samples. One sample was incubated with 10 mM MTSES, one was incubated with 10 mM NEM, and the 2 remaining samples were incubated in the absence of thiol reactive reagent (replaced with an equal volume of water). The samples were incubated at room temperature in the dark for 1 hour. The treated cells were harvested by centrifugation, washed with PBS to remove excess MTSES and NEM, resuspended in lysis buffer (15 mM Tris pH 7.6, 1% (w/v) SDS, 6.2 M urea), 6.25 mM mPEG5K, DNase and protease inhibitors, and incubated at room temperature for 1 hour. SDS-PAGE sample buffer was added to the samples, which were then separated using SDS-PAGE and visualized using Wester blotting with an anti-his tag antibody (Invitrogen).

### Copper phenanthroline-based crosslinking

Cell samples containing overexpressed YqjA variants were prepared as described above for the SCAM assay. Once prepared, the samples were incubated at room temperature in the presence of 1 mM dithiothreitol (DTT), or 0, 0.1, 0.3 or 0.6 mM Copper phenanthroline (CuPhen). Crosslinking was quenched by addition of 100 mM Methyl-methanethiosulfonate (MMTS), which also prevented further disulfide formation upon denaturation of the protein by alkylating any remaining free cysteines. SDS-PAGE sample buffer, DNase and protease inhibitors were added to the samples, which were then separated using SDS-PAGE and visualized using Wester blotting with an anti-his tag antibody (Invitrogen).

### Topological and hydropathy profile analysis

We generated our hydropathy profile of YqjA using ExPASy ProtScale tool (https://web.expasy.org/protscale) using the Kyte-Doolittle hydropathy scale with a residue window of 11. Topological analysis was performed using the TOPCONS tool (https://topcons.cbr.su.se/pred/) and TMHMM (http://www.cbs.dtu.dk/services/TMHMM/) using default settings ^23,24^.

### Generation of structural model using evolutionary covariance

Structural models for YqjA were generated using EVfold^25^ using the default parameters. High quality modelling data were generated for various bitscore thresholds (0.1, 0.3, 0.5 and 0.7). We selected the model generated from the 0.7 threshold dataset due to it having the most sequence coverage and because it produced the highest number of feasible models. Models were visualized using PyMol, which was also used to generate the structural images.

## Results

### Consensus topological analysis suggest YqjA contains 5 transmembrane helices

To provide preliminary information on the topological arrangement of YqjA, we determined a hydropathy plot of YqjA, which provides an indication of the level of hydrophobicity in the primary sequence. Analysis of the YqjA hydropathy plot reveals 6 distinctly hydrophobic regions that likely relate to transmembrane spanning regions (Fig. 1A). To take this analysis further, we used TOPCONS and TMHMM to generate a consensus predicted topology for YqjA^23,24^. TOPCONS compares the topology predictions from 5 separate topology prediction algorithms (OPTOPUS, Philius, Polyphobius, SCAMPI and SPOCTOPUS) and produces a consensus predicted topology. Comparison of the outputs from different prediction algorithms revealed substantial variation in the number of predicted transmembrane regions and the location of N-terminus; 4 out of the 5 algorithms predicted 5 TMs with a Nout/Cin orientation, although there was variation in the location of the TMs (Fig. 1B). Philius predicted 6 TMs with a Nin/Cin orientation, which was also the same prediction made by TMHMM (Fig. 1B). On balance of all the predictions generated, the consensus topology model for YqjA is 5 TM with Nout/Cin (Fig. 1C).

**Figure 1.**
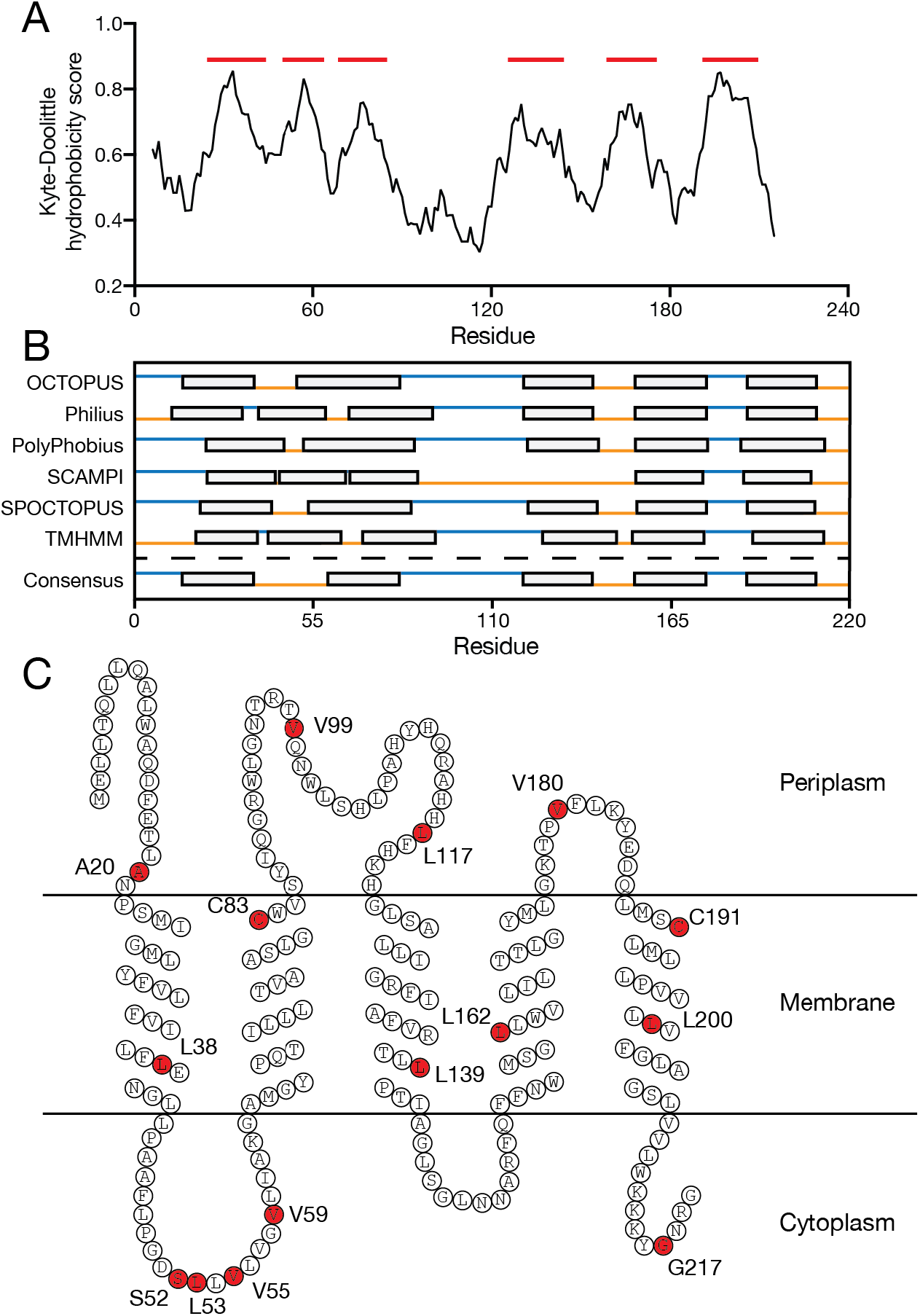
Hydropathy profile analysis and 2D topological predictions of YqjA. **A)** Hydropathy profile analysis of YqjA based on the Kyte-Doolittle hydropathy scale and a window of 11 residues. **B)** Comparison of the topological prediction generated by TOPCONS and TMHMM. Grey boxes represent TM helices, blue lines indicate periplasmic regions and orange lines indicate cytoplasmic regions. **C)** Schematic of the consensus topology of YqjA. Red coloured amino acids indicate those that we mutated to cysteine for SCAM analysis.

### SCAM analysis of YqjA reveals a topological map incongruent with a 5 TM model

To experimentally probe the membrane topology of YqjA, we performed substituted cysteine accessibility method (SCAM). While there are many variants of the SCAM approach^26^, the basic premise is that single cysteines are individually introduced into an integral membrane protein, and their location in relation to the membrane (periplasmic or cytoplasmic) is assessed by the accessibility of each cysteine to thiol-reactive reagents that are either permeable or impermeable to the intact membrane. Here, we employed an approach similar to SCAM approaches used previously^27–29^.

To perform SCAM on YqjA, we introduced single cysteine residues individually into a cysteine-free variant of YqjA to produce a library of single cysteine mutants. Each member of the mutant library was then expressed independently in *E. coli* from a plasmid in-frame with a C-terminal histidine tag allowing for detection of expressed YqjA in whole cell extracts via Western blotting. We harvested the cells expressing the cysteine variants and incubated samples with either 2-sulfonatoethyl methanesulfonate (MTSES), which is a cysteine-reactive reagent impermeable to the inner membrane (but can traverse the outer membrane), or N-ethylmaleimide (NEM), which is permeable to both *E. coli* membranes. Thus, MTSES would only conjugate to cysteines accessible to the periplasm, whereas NEM could react with cysteines accessible to both the periplasm and cytoplasm. Cysteine residues buried in the protein core or in the middle of a transmembrane region would likely react with neither MTSES nor NEM. Conjugation of the introduced cysteines to MTSES or NEM would protect that position from further thiol-specific reactions. Thus, the level of protection afforded by MTSES and NEM to further reaction is an indicator of its position relative to the membrane. To assess the level of thiol protection, we solubilised the membrane and denatured the protein using SDS, and incubated the sample with methoxypolyethylene glycol maleimide (mPEG5K), which reacts with free cysteines to add 5 kDa mass to the protein that would be separable from the unmodified protein using SDS-PAGE. We then visualized YqjA using Western blotting with an anti-his tag antibody to assess the level of YqjA PEGylation.

To perform SCAM analysis of YqjA, we first needed to produce a cysteine-free version of YqjA by mutating the two native cysteines, C83 and C191, to serine (hereafter referred to as YqjAcysless). To test that the YqjAcysless was fully functional, we expressed the mutated gene from an arabinose-inducible plasmid and assessed its ability to restore growth in the double *dedA* deletion *E. coli* strain, BW25113ΔyqjAΔyghB (BW25113ΔΔ, hereafter), which is unable to grow at elevated temperatures (the same phenotype seen for the well characterized strain BC202, which has the same double *dedA* deletion in an *E. coli* W3110 background^5^). Expression of both wildtype YqjA (YqjAwt) and YqjAcysless restored growth to BW25113ΔΔ at 44°C, whereas expression of an unrelated integral membrane protein, the aspartate transporter Glt_Ph_, was unable to restore growth at this temperature, demonstrating that YqjAcysless was functional and folded (Fig 2A). Using YqjAcysless as a background, we generated a panel of 15 single cysteine YqjA mutants with cysteines distributed throughout the amino acid sequence of YqjA (Fig. 1C). We established that all 15 single cysteine YqjA mutants were functional as demonstrated by their ability to restore growth to BW25113ΔΔ at elevated temperatures (Fig. 2A), giving us confidence that they would accurately report on the topology of the fully folded protein.

**Figure 2.**
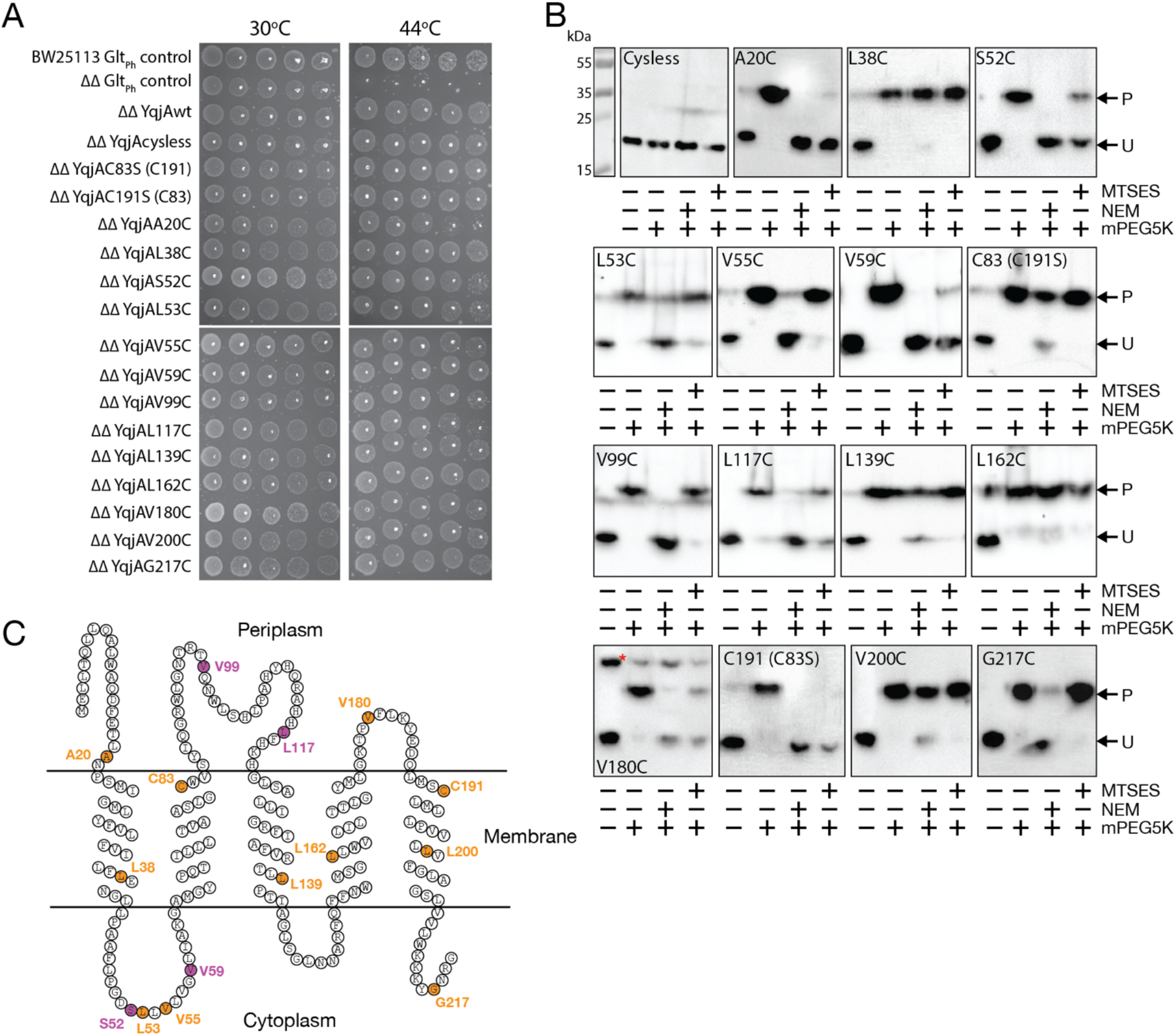
SCAM analysis of YqjA. **A)** Phenotypic rescue assay in which BW25113 wildtype or BW25113ΔΔ cells (ΔΔ) harbouring plasmids expressing *yqjA* variants or a control plasmid (Glt_Ph_ control) were grown on LB agar supplemented with 0.001% L-arabinose at 30°C and 44°C. This experiment was performed 3 times with the same result. **B)** Western blot analysis of whole cell lysates from cells harbouring plasmids expressing cysteine free (“Cysless”) or single cysteine variants of *yqjA* with and without treatment with NEM, MTSES and mPEG5K. PEGylated (P) and unmodified (U) protein are indicated with arrows. The data presented is representative of at least 2 separate experiments for each mutant (number of replicates shown in Supplementary Table 1). **C)** 2D topological map of YqjA with the positions of the cysteines colour coded to indicate agreement between the 2D map and the SCAM analysis; orange indicates agreement between 2D map and SCAM data, purple indicates disagreement.

Each of the single cysteine variants was expressed in *E. coli* and subjected to the SCAM procedure described previously. Treatment of each of the 15 cysteine mutants with mPEG5K alone resulted in a single higher molecular weight band demonstrating that each protein contained a single cysteine, which was able to react with cysteine reactive reagents (Fig. 2B). Based on the assumption that MTSES would only protect against PEGylation for cysteines exposed to the periplasm, and NEM would protect cysteines exposed to both the periplasm and cytoplasm, our SCAM data suggested that A20C, S52C, V59C and V180C were exposed to the periplasm, L53C, V55C, V99C, L117C and G217C are exposed to the cytoplasm, and L38C, C83, L139C, L162C, C191, and V200C are all buried in the membrane/protein core (Fig. 2B).

Mapping these experimentally-defined accessibility measurements onto the 2D topology models revealed that while most of the locations determined by SCAM matched the predicted topology, 4 positions were the complete opposite; V99C and L117C were predicted to be periplasmic, but were located in the cytoplasm by SCAM, whereas S52C and V59C were predicted to be cytoplasmic but were located in the periplasm according to our SCAM data (Fig. 2C). These data suggest that YqjA adopts a substantially different arrangement than that represented in the 2D models.

### V180 likely forms part of an oligomeric interface

While performing our SCAM analysis, we noticed that unlike all the other single cysteine mutants, V180C had a substantial band in the Western blot at approximately 50 kDa. This ~50 kDa band was present in all of the samples for V180C, but was most prominent in the sample that was incubated in absence of any cysteine labelling reagent (Fig. 2B). The molecular weight of this band corresponds approximately with that of dimeric YqjA, which we reasoned was likely stabilized by inter-protomer disulfide formation between V180C residues, as has been seen previously for C191 in YqjA^30^. To investigate this possibility further, we incubated YqjAV180C-expressing cells with increasing concentrations of the oxidizing agent copper phenanthroline (CuPhen) and observed a CuPhen concentration-dependent increase in the ~50 kDa band intensity with a concurrent decrease in the intensity of the band corresponding to the YqjA monomer (Fig. 3A). In addition, we observed no higher molecular weight band in the presence of reducing agent dithiothreitol (DTT), nor when YqjAcysless was incubated with the same range of CuPhen concentrations (Fig. 3A). To investigate whether this phenomenon was specific to V180C and to rule out the possibility that we are observing crosslinking of the cysteines after the protein has been denatured for SDS-PAGE analysis, we incubated YqjAA20C-expressing cells with the same CuPhen concentrations; A20C is located on the periplasmic side of the YqjA but in a different region of the protein to V180C. We observed no higher molecular weight band in the absence of CuPhen for YqjAA20C, and only very minimal apparent crosslinking at the highest CuPhen concentration (Fig. 3A). Taken together, our data suggest that V180C from two YqjAs are able to form an intermolecular disulfide, which is formed when the protein is folded in the membrane. Due to the close proximity required to form a disulfide, these data suggest that YqjA is an oligomer (although the oligomeric state is not possible to glean from these data), the region containing V180C likely forms an oligomeric interface, and due to V180C being located on the periplasmic side of the protein, this conclusion suggests that the proteins involved in disulfide bond formation are arranged in a parallel fashion, contrary to previous suggestions that DedA form an antiparallel dual topological arrangement (Fig. 3B)^17^.

**Figure 3.**
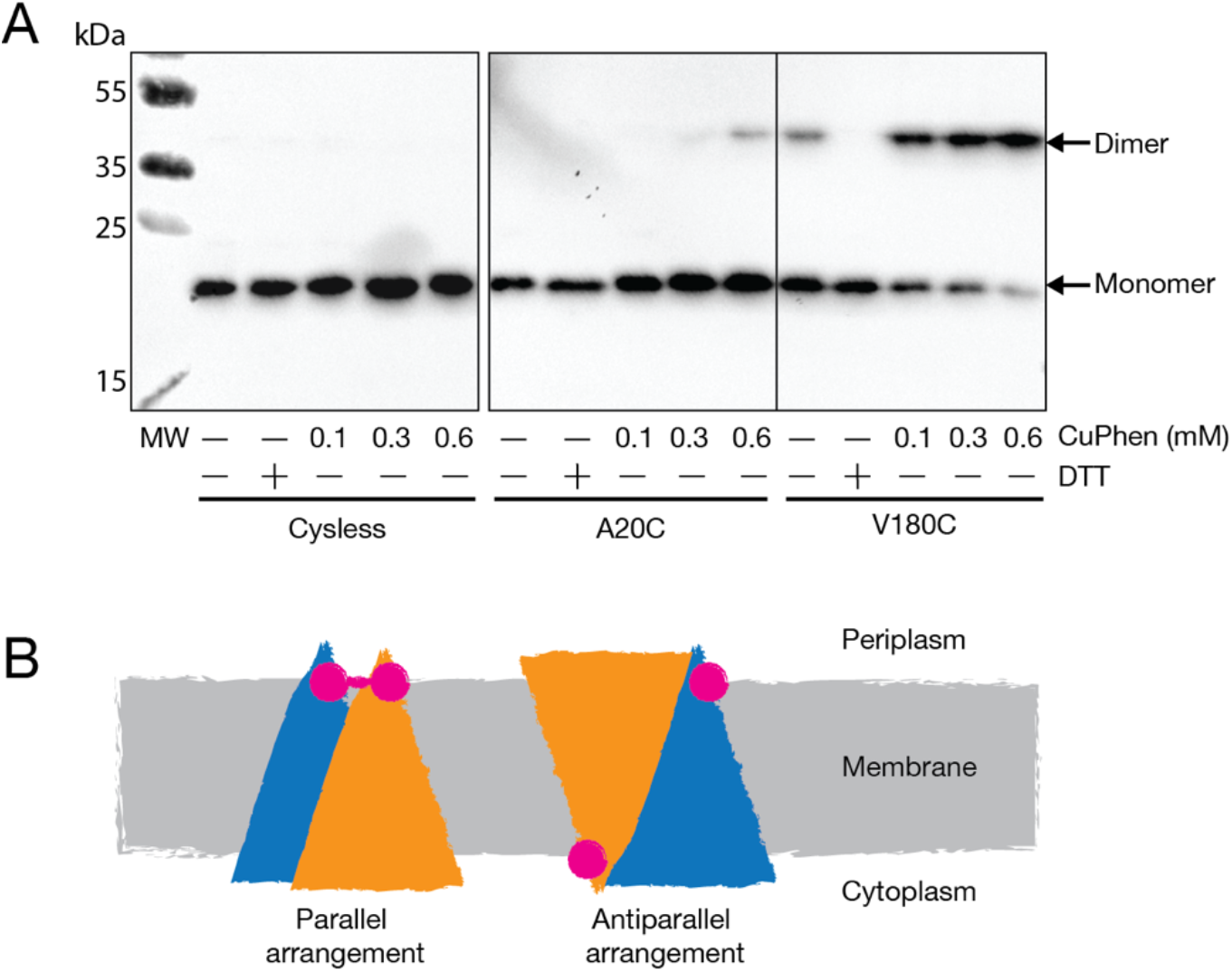
Disulfide crosslinking of YqjAV180C. **A)** Western blot analysis of whole cell lysates generated from cells harbouring plasmids expressing variants for *yqjA* treated with reducing agent DTT or increasing concentrations of the oxidizing agent CuPhen. This experiment was performed on two separate occasions with the same result. **B)** Schematic showing the relationship between protomer orientation in parallel (left) and antiparallel (right) arrangements. The protomers in homooligomeric YqjA are coloured orange and blue for clarity, and the approximate position of V180C is represented by a pink circle. In the parallel arrangement, the cysteines can form a disulfide (represented by the pink line between cysteines), whereas, in the antiparallel arrangement, a disulfide cannot form.

### SCAM analysis supports a model for YqjA based on evolutionary covariance analysis

*Ab initio* models of members of the DedA superfamily have been generated using evolutionary covariance analysis using trRosetta, which suggest DedA proteins form an α-helical bundle and contain 2 re-entrant hairpin loops^2,19^. To generate a 3D model of YqjA on which to map our SCAM data, we also used evolutionary covariance analysis, but with EVfold, which has also been used previously for a DedA family member, but distantly related eukaryotic homologue of YqjA^2^.

EVfold analysis of YqjA produced high quality modelling data (according to the EVfold output) for datasets with bitscores ranging from 0.1 (48199 sequences) to 0.7 (3917 sequences). For the generation of the working model for our analysis, we selected the model generated using the dataset with a bitscore of 0.7 because it provided the best protein sequence coverage and the majority of the models it produced (29 out of the top 30 scoring models) modelled the C-terminal helix to produce a sensible arrangement (overlay of the top 30 scoring models is shown in SI fig. 1A). One model generated using the 0.7 bitscore dataset arranged the C-terminal helix in a position where it would be inserted into the core of the bilayer, which we do not consider feasible, and not compatible with our experimental data (SI fig. 1A). While the highest scoring model generated using the greatest evolutionary depth dataset (bitscore of 0.1) produced a 3D model similar to 0.7 bitscore dataset, the majority of the other high scoring models generated using this dataset mishandled the C-terminal helix to produce a series of what we consider to be unfeasible structures that are also inconsistent with our experimental data (overlay of the top 30 scoring models from the 0.1 bitscore dataset is shown in SI fig. 1B). This mishandling of the C-terminal helix is likely due to this region of YqjA being involved in the formation of an oligomeric interface.

As with previous *ab initio* models generated for DedA superfamily members^2,19^, the structural model for YqjA generated using EVfold produces an α-helical bundle consisting of 3 membrane spanning helices, 2 re-entrant hairpin loops and a short α helix perpendicular to the cytoplasmic side of the membrane (Fig. 4 A and B). While the evolutionary co-variance approach cannot assign membrane orientation, all of the *in silico* prediction software predict a cytoplasmic N-terminus (Fig 1B), which also matches our experimental SCAM data. The tips of the two predicted re-entrant hairpins, which we have named HPin and HPout, are predicted to meet in approximately the centre of the membrane (Fig. 4C). Mapping on the known functionally essential residues, E39, D51, R130 and R136^6,30,31^, we note with interest that they are clustered at the interface of these 2 predicted hairpins, suggesting that this region constitutes a crucial active/binding site, and providing support for this structural model (Fig. 4D).

**Figure 4.**
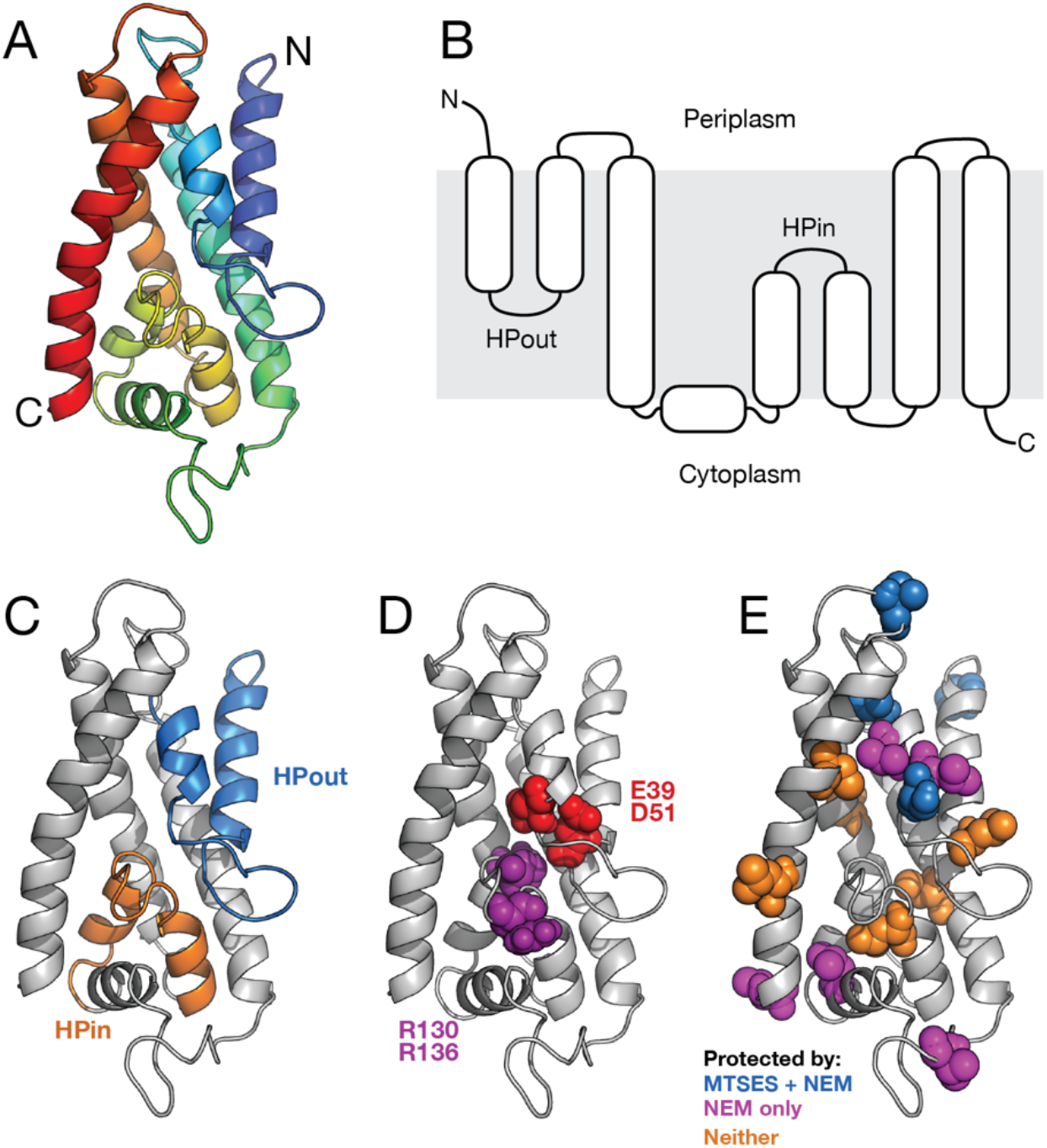
*Ab initio* model of YqjA using co-evolutionary variance. **A)** Cartoon representation of the 3D structure of YqjA generated using EVfold analysis. The main chain is represented as a ribbon, rainbow coloured with N-terminus blue and the C-terminus red. **B)** 2D representation of the 3D model in A). **C)** YqjA structural model with the predicted hairpins highlighted. **D)** YqjA structural model with the clustered functionally essential residues highlighted. **E)** YqjA structural model with our SCAM data mapped on. Blue positions were protected by both MTSES and NEM, orange positions were protected by neither MTSES nor NEM, and magenta residues were protected by only NEM. P22 is highlighted in place of A20 and V209 is highlighted in place of G217 because these positions were not covered in the model.

Mapping our experimental SCAM data onto the EVfold structural model, we find that the cysteine protection we observed by MTSES and NEM is structurally rationalized for 13 out of the 15 positions tested (Fig. 4E). A20C, S52C, V59C, and V180C, which were all protected by the membrane impermeable MTSES, suggesting they are accessible to the periplasmic side of the membrane, are clearly positioned on the periplasmic side of the protein with 3 of the positions located in loop regions (Fig. 4E, blue residues). L38C, C83, C191, L139C, L162C, and L200C, which were protected by neither MTSES nor NEM, suggesting they are embedded in the protein/membrane core, form a band around the centre of the protein which is likely membrane embedded, explaining the lack of accessibility (Fig. 4E, orange residues). V99C, L117C and G217C, which were only protected by NEM, suggesting a cytoplasmic location, are positioned in cytoplasmic loops in the structural model.

Residues L53C and V55C are predicted to be positioned in the arm of HPout on the periplasmic side of the protein, but are only protectable by NEM (Fig 4E and 2B). While this may make the structural model and SCAM data seem incongruent, the exclusive accessibility of L53C and V55C to NEM can be explained by one of two possibilities; NEM, which is relatively hydrophobic compared to the negatively charged MTSES, is able to penetrate deeper into hydrophobic pockets on the periplasmic side of YqjA; or, conformational changes occur in the re-entrant hairpin loops, as seen for hairpin-containing secondary active transporters^32–35^, expose that region of HPout to the cytoplasmic solution. The fact that S52C, which is obviously proximal to L53C and V55C due to its position in the primary sequence, is protectable by MTSES, which *cannot* penetrate the bilayer, very strongly supports the positioning of this region on the periplasmic side of the protein. Therefore, our SCAM data support the EVfold model, but more information on the structure and dynamics of YqjA is required to fully rationalize all of the SCAM data collected. To provide further support for the EVfold model, we also obtained the structural model for YqjA from the Alphafold Protein Structural Database^36^. The Alphafold model of YqjA has the same overall topology as the EVfold model and contains the two re-entrant hairpin loops (SI fig. 2). In addition, the accessibility of each cysteine from our SCAM data can be rationalised similarly in both models. The primary difference between the two YqjA models is the positioning of the C-terminal helix, which overlays HPin in the Alphafold model but is adjacent to HPout in the EVfold model (SI fig. 2B). However, with the C-terminal helix likely involved in homo-oligomer interface formation, it is a problematic region to model using evolutionary coupling due to the difficulty in differentiating between intra- and inter-protomer residue coupling.

## Discussion

In this study, we have performed a cysteine accessibility scan on the most comprehensively characterized DedA protein, YqjA from *E. coli* and determined the relative position of 15 individually substituted cysteine residues to the membrane. By comparing our experimental SCAM data to a 2D topological model obtained using traditional approaches, we find our experimental data in disagreement with the 2D topology map in several positions. However, our SCAM data is in good agreement with a structural model generated using evolutionary covariance. This is the first experimental verification of a 3D structural model of a bacterial DedA family member. The structural model of YqjA predicts the presence of 2 re-entrant hairpin loops, the tips of which converge to bring 4 functionally relevant amino acids into close proximity, likely forming the binding or active site of the protein. This experimentally tested model supports a recent model of YqjA generated using a similar modelling approach^19^, and with a structural model of a distantly related eukaryotic DedA superfamily member^2^. Our SCAM analysis of YqjA also suggests that V180 is able to form interprotomer disulfide bonds, which strongly suggests that YqjA is an oligomer, and due to the position of V180 on the periplasmic side of the membrane, these data indicate that the oligomer forms a parallel arrangement in the membrane.

### Re-entrant hairpins likely play a fundamental role in YqjA function

Understanding the structure of the DedA superfamily is key to understanding their function and mechanism. There are currently no high resolution structures for any DedA superfamily member, and analysis with the Phyre2 server suggests that there are currently no related, structurally characterised proteins on which one could model a DedA structure^37^. The recent advances in protein modelling via evolutionary covariance combined with structural verification via SCAM analysis is a powerful and highly accessible approach to determining structural characteristics of integral membrane proteins of unknown structure. There have now been three modeling studies of the DedA superfamily using evolutionary covariance, and while some of the details of helical placement may not be in complete agreement between the models, the presence of 2 re-entrant hairpins that converge in the centre of the protein is common among the models^2,19^. Re-entrant hairpin loops are regions of integral membrane proteins that dip into and then exit the bilayer on the same side of the membrane. This structural feature has been identified in several integral membrane protein structures, most commonly associated with ion-driven secondary active transporters, including members of the excitatory amino acid transporter (EAAT) family and their prokaryotic homologue, Glt_Ph_^38,39^, members of the divalent anion sodium symporter (DASS) family^40–42^, and VcCNT^43^, and without exception, the tips of these hairpin loops form a crucial site for substrate interactions. Re-entrant loops are also thought to be involved in gating access to the binding site and undergo conformational changes by which they control ingress and egress of the substrate(s) to and from the binding site^32,33,35,44^. Four functionally essential residues have been identified in YqjA; E39, D51, R130 and R136^6,30,31^, and while the exact role these residues play in the function of YqjA is unknown, they coalesce at the tips of the re-entrant hairpins, demonstrating the importance of this structural motif in YqjA. While direct functional measurements are yet to be made for any member of the DedA superfamily, there is strong circumstantial evidence that at least one of the functions of YqjA is as a monovalent cation/H^+^ exchanger. For example, YqjA is required for growth at high pH^8^, and alkalinotolerance mechanisms often involve Na^+^ or K^+^/H^+^ exchangers, e.g. NhaA and MdtM^45–47^. In addition, YqjA contains membrane embedded acidic residues that are essential for its ability to rescue growth defects caused by disruption of *yqjA* and *yghB* genes in *E. coli*^10^. Furthermore, the growth defects observed in the *E. coli* strains in which *yqjA* and *yghB* are disrupted can be rescued by lowering the external pH to artificially bolster the proton motive force (PMF), by increasing the external monovalent cation concentration, or by overexpressing *mdfA* which encodes an multidrug efflux pump that also has Na^+^/H^+^ activity^10^. Due to the positioning of the well-conserved, functionally important residues at the tips of the re-entrant hairpins, it is likely that this is the substrate binding site with the binding and release of protons by E39 and D51 a central part of the mechanism. However, more structural information is required to resolve the details of this protein region, identify the exact position of the binding site, identify the ligand(s), and to assess the conformational dynamics that may be required for its mechanism.

### YqjA likely forms a homooligomer

Our SCAM data show that the YqjAV180C variant produced a higher-than-expected molecular weight band on the Western blot (Fig. 2B), which we demonstrated is likely due to disulfide formation between cysteines in the same position in two protomers, suggesting that YqjA forms an oligomer. The fact that only one of the cysteine mutants gave a dimeric species indicates that this dimerization is likely due to close proximity of the V180 residues. It has previously been suggested that *E. coli* YqjA is able to form an oligomer due to the observation that the native cysteine residue C191 and a cysteine substituted for L195 are capable of forming an intermolecular disulfide with the equivalent residue in a vicinal protomer, similar to our observation of V180C^30^. Combined, these observations suggest that a large part of the predicted C-terminal helical region of YqjA forms an oligomeric interface. However, the actual oligomeric state of YqjA and the DedA superfamily in general is still an open question. Based on the previously observed disulfide bond formation between C191 and L195C in YqjA, it has been asserted that the protein forms a dimer. However, as disulfides would only ever form between a maximum of 2 cysteines, a dimeric band in the membrane would also be observed if YqjA formed a higher oligomeric state, for example, a trimer or a tetramer. Therefore, more detailed biochemical analyses of purified DedA superfamily members is required before a definitive oligomeric state can be concluded.

### All YqjA proteins are oriented identically in the membrane

Based on hydropathy profile alignments using the AlignMe program, it was hypothesized that the DedA family shares a common ancestor with LeuT-fold transporters^48^. The leuT fold core is composed of an inverted structural repeat of 5 transmembrane helices that evolved via duplication and fusion of an ancestral gene^49,50^. However, for the repeats to be inverted, the ancestral gene would need to produce a protein that was able to reside in the membrane in a mixed topology. Due the marked similarity between the hydropathy profiles of certain DedA proteins and one repeat of selected LeuT fold proteins, it was suggested that the DedA family could represent a “half-module” of the LeuT fold^17^. Our SCAM data demonstrate that YqjA resides in the membrane in a single orientation. Firstly, if YqjA had a mixed topology in the membrane then we would expect to observe incomplete protection by MTSES, which is impermeable to the membrane, for all positions predicted to be accessible to the periplasm or cytoplasm. However, in all cases, MTSES fully protects cysteines accessible to the periplasm, but has no effect on cysteines located in the cytoplasm. Secondly, V180C is able to form an interprotomer disulfide bond, and as this position is predicted, and shown experimentally, to be present in a periplasmic loop, this observation strongly suggests that the two crosslinked protomers are adopting the same orientation in the membrane. Taken together, these data demonstrate that YqjA is present in the membrane in a single orientation. However, what is true for *E. coli* YqjA may not be the case for other members of the DedA superfamily as there are clearly different functional DedA groups even within the same organism^3^, and there is strong evidence that some DedA proteins have ambiguous charge bias^17^, which could allow them to be dual topology. Therefore, it is possible that some DedA proteins adopt a dual topology, but further work is required to resolve this issue.

Here, we have combined evolutionary covariance modelling with cysteine labelling to generate and experimentally validate a structural model of the archetypal DedA protein, YqjA from *E. coli*. This work provides insight into the architecture of bacterial DedA proteins and will aid in the delineation of the function and mechanism of this widespread, physiologically important protein family.

## Author contributions

HS, RU and BS performed experiments. AM supervised work and performed experiments. CM supervised the experiments, analysed data, conceived project and provided resources.

## Conflicts of interest

The authors report no conflicts of interest.

## Acknowledgements

We thank Lucy Forrest, Vanessa Leone, Ben Goult and Matthew Batson for helpful suggestions. We thank Tracy Palmer for the kind gift of several *E. coli* knockout strains. This work was supported by the Royal Society Research grant (grant no. RG170265) awarded to CM.

## Supplementary Information

**Supplementary table 1.**
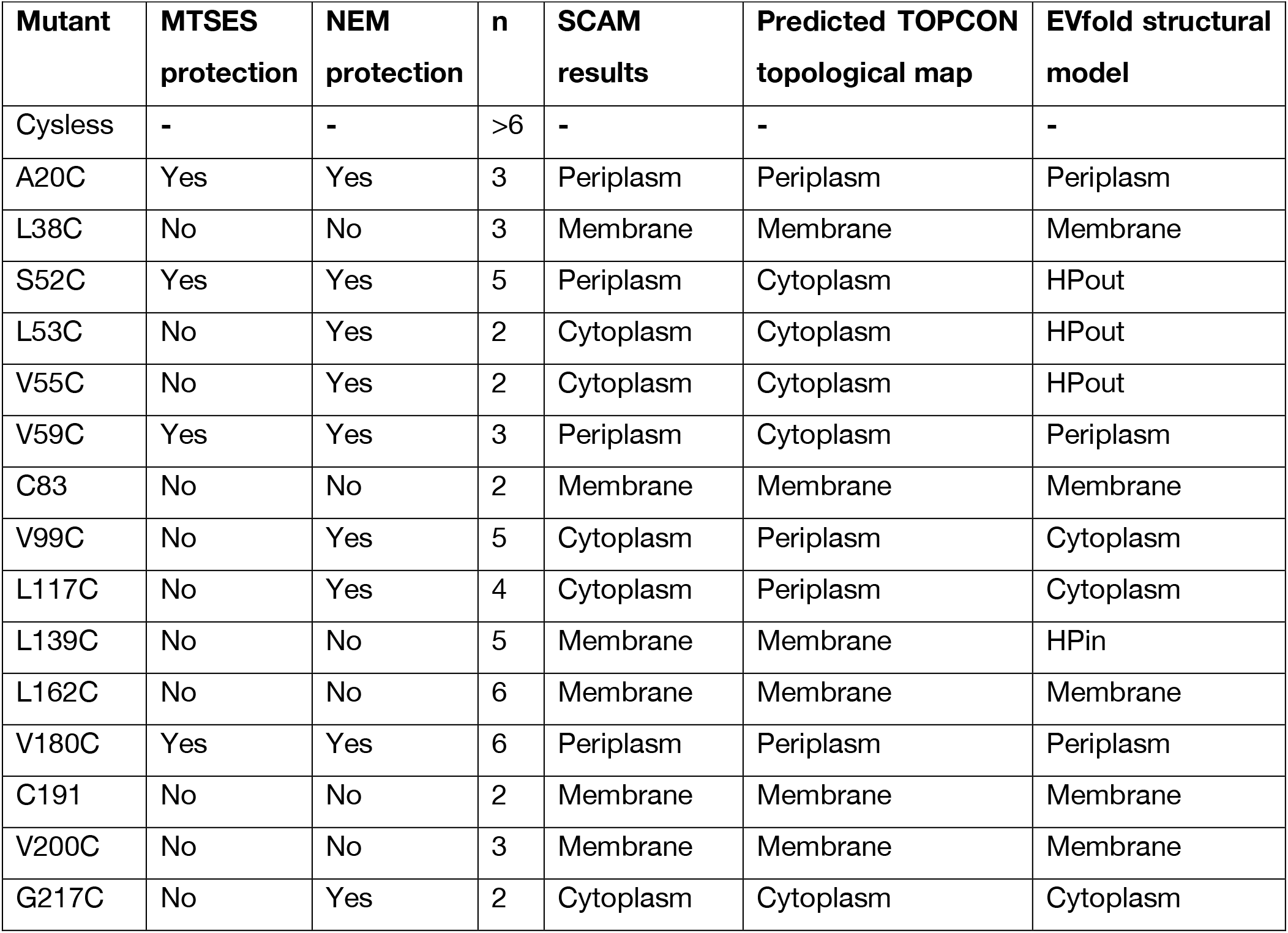
Summary of SCAM results for the panel of single cysteine mutants, the number of replicates tested for each position, and the predicted location of each cysteine according to the 2D and 3D models.

**Supplementary figure 1.**
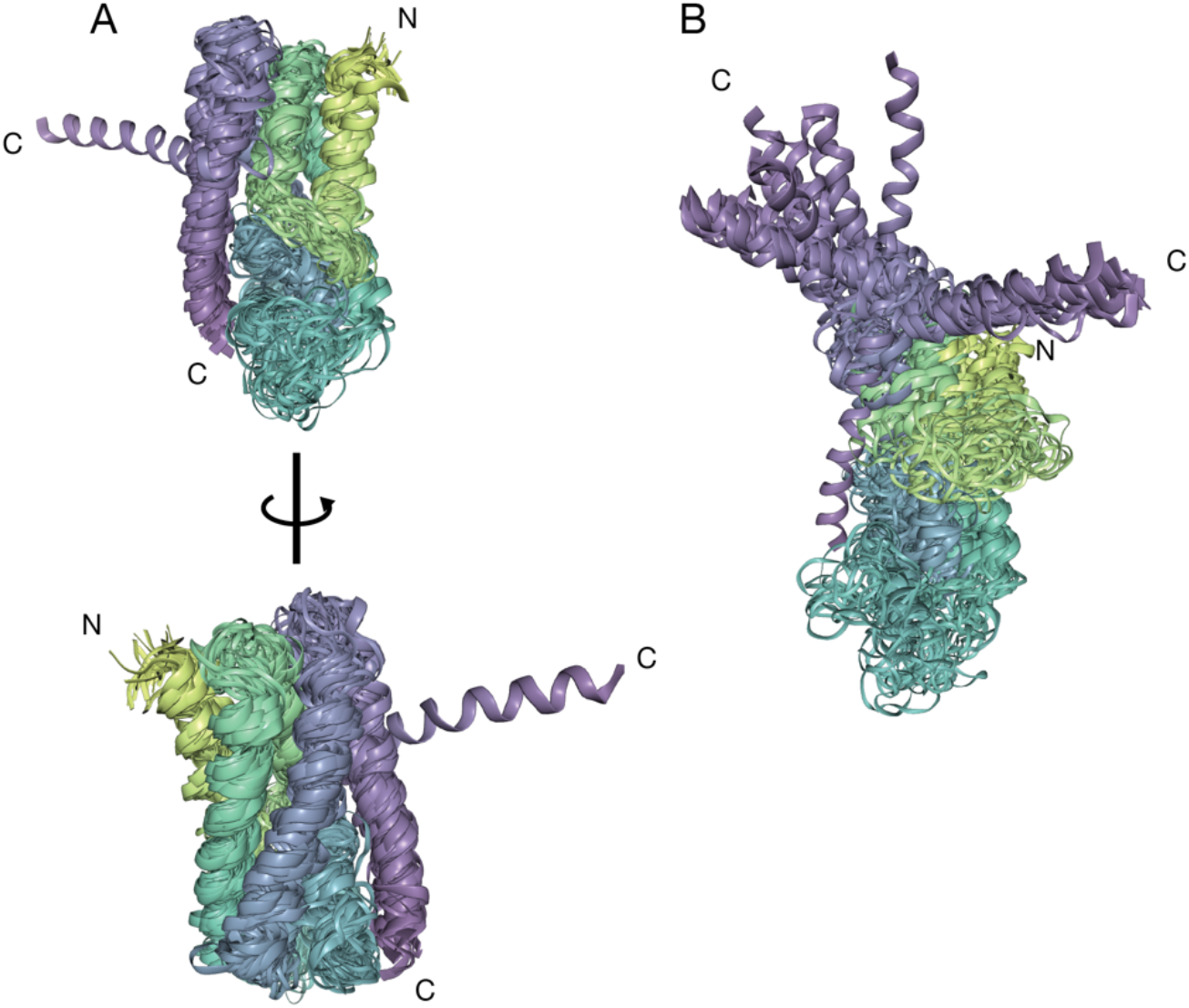
Overlay of models generated by EVfold. **A)** Overlay of the top 30 scoring models form the 0.7 bitscore dataset. 29 out of 30 superimpose well and form a tight α-helical bundle. The remaining model superimposes with the other models for the majority of the structure, however, the C-terminal α-helix (purple helix) is modelled at an angle incongruent with our SCAM data. **B)** Overlay of the top 30 models generated using the 0.1 bitscore dataset. Only 1 model out of 30 modelled the C-terminal helix (purple helix) in an arrangement in agreement with our SCAM analysis, the rest modelled this helix in a variety of unlikely positions, none of which agree with our SCAM data.

**Supplementary figure 2.**
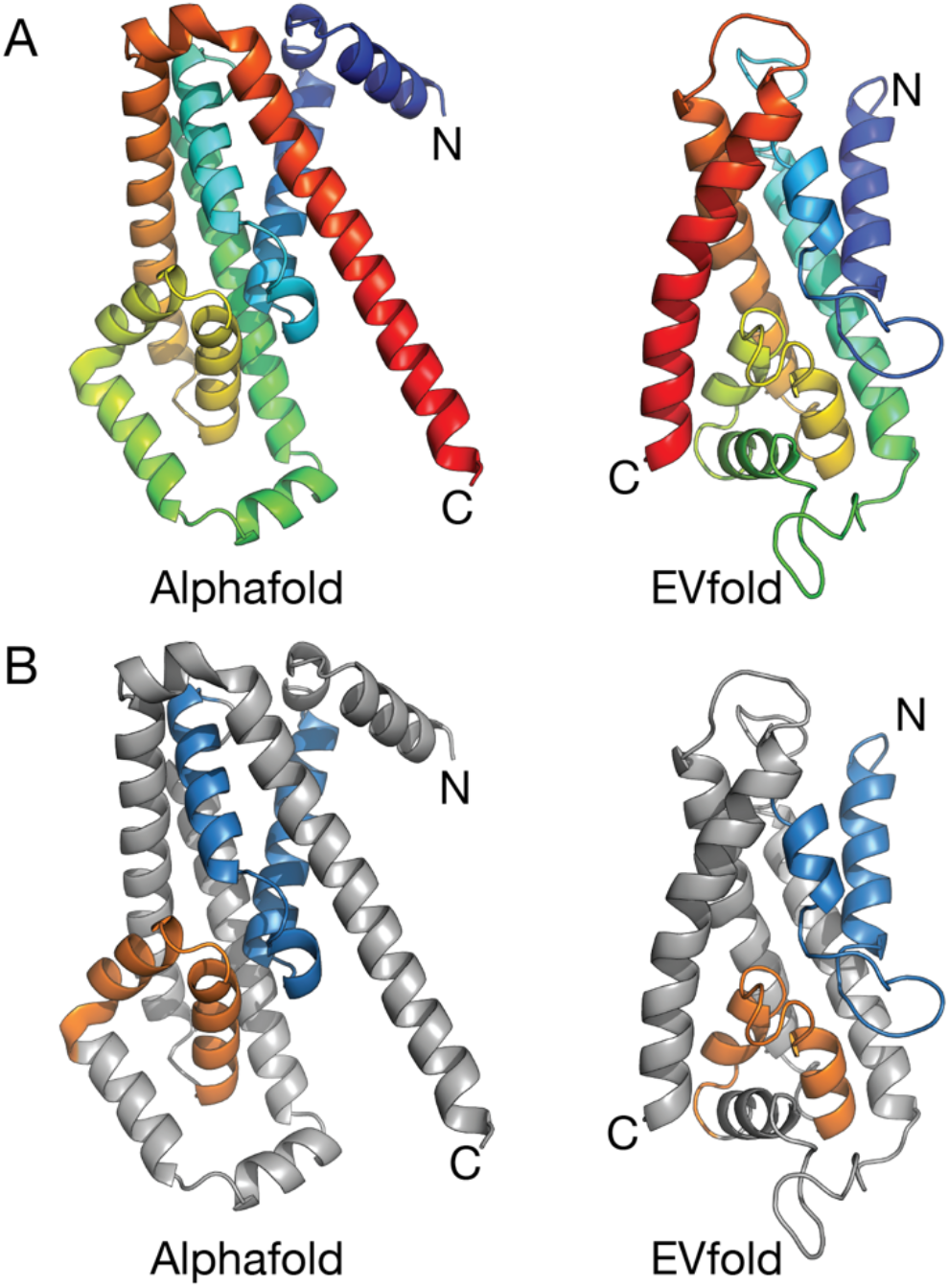
Comparison of YqjA models generated by EVfold and Alphafold. **A)** Rainbow-coloured ribbon representation of the Alphafold (left) and EVfold (right) models of YqjA oriented according to the hairpin loop positioning. Note the main difference between the models being the positioning of the C-terminal helix coloured in red in both cases. **B)** The same models as in A), but with the predicted re-entrant hairpin loops highlighted (HPout in blue, HPin in orange).

## Notes

### Competing Interest Statement

The authors have declared no competing interest.

